# Layer connective fields from ultrafast resting-state fMRI differentiate feedforward from feedback signaling

**DOI:** 10.1101/2025.02.23.639720

**Authors:** Joana Carvalho, Francisca F. Fernandes, Mafalda Valente, Koen V. Haak, Noam Shemesh

## Abstract

Deciphering the directionality of information flow in cortical circuits is essential for understanding brain dynamics, learning, and neuroplasticity after injury. However, current non-invasive methods cannot distinguish feedforward (FF) from feedback (FB) signals across entire networks, including deep brain regions. Here, we present a novel approach combining ultrafast fMRI with a Layer-based Connective Field (lCF) model to disentangle FF from FB signaling. Our findings reveal that lCF size, an indicator of spatial information integration, differentiates FF and FB activity through distinct layer-specific connectivity patterns during spontaneous activity, challenging the notion that FF signals are solely stimulus-driven. FF connectivity follows an inverted U-shape, peaking in layer IV, while FB exhibits a U-shaped pattern, with peaks in layers I and VI. These profiles generalize across sensory pathways (visual, somatosensory, and motor) and reveal injury-induced network reorganization, such as LGN bypassing V1 to provide direct FF input to higher visual areas.

## 1. Introduction

Sensory processing depends on hierarchical and reciprocal communication across brain regions^1–4^, with signals flowing dynamically between early sensory areas and higher cortical areas^5–7^ in multiple species^8,9^. This bidirectional information flow allows early sensory areas to relay encoded sensory information to higher cortical regions for further processing, which is then fed back to refine or modulate the initial processing in the early regions^10–12^. Feedforward (FF) processing entails a bottom-up flow of neural signals, where simple receptive fields in primary sensory areas build up to more complex representations in higher-order areas^2^. In contrast, feedback (FB) signalling introduces top-down influences, integrating elements such as attention^12,13^, prediction^14^, and contextual cues to shape perception in a meaningful, adaptive way^15^.

Differentiating the directionality of information flow in cortical circuits is key to understanding neural computations underlying learning, perception, and neuroplasticity^16–19^. However, dissociating FF from FB signals remains challenging due to their concurrence and their distribution in wide networks in the brain. So far, studying FF and FB signals relies on invasive methods like electrophysiology, calcium imaging, photometry^20^, and optogenetics, which provide high specificity but are limited to subsets of neurons, potentially missing pathway-wide mechanisms^21–23^. In humans, electrocorticography has been used but remains invasive and localized^11^. Non-invasive techniques like MEG and EEG offer good temporal resolution but lacking the spatial precision and depth required to investigate local circuits, such as those within cortical columns^24,25^, or more broadly distributed systems. Additionally, signal overlap from neighboring areas complicates FF and FB segregation in these techniques.

Functional MRI provides a critical view into brain wide signal processing ^26–28^. Ultrahigh-field MRI provides noninvasive access to high spatial resolution^29,30^ and can enable the segregation of cortical layer signals through neurovascular coupling^31–33^, offering insights into FF and FB processing^29,34-,36^. Previous research has often relied on manipulating experimental paradigms to isolate FF and FB signals, such as occlusion paradigms^37^, visual illusions^38^ and contrasting visual perception with imagery^39^. FF and FB signals can be distinguished based on their cortical layer of origin and reception^40^. FF signals typically originate in middle cortical layers^41^, while FB signals show variability, arriving in deep^42,43^, superficial^37,41^, or both layers^44,45^. Despite these advances, noninvasively distinguishing FF from FB signals has not been achieved, due to their spatiotemporal overlap.

The temporal dynamics of FF and FB signals have been largely overlooked in fMRI due to conventional methods’ limited resolution and slow and variable hemodynamic coupling. Notably, FF and FB signals exhibit distinct temporal characteristics: FF operates at higher frequencies (30-50 Hz) and occurs shortly after stimulus onset, while FB is slower (3-6 Hz)^6^ and more prevalent during spontaneous activity, making their interaction with neurovascular coupling interesting to target^46^. Advances in preclinical high-field MRI now allow sub-millimeter resolution (∼150 µm) and rapid sampling (∼30-50 ms)^47–49^, enabling the accurate mapping of stimulus-evoked neural input mapping^48,50^. Critical developments in ultrafast fMRI acquisitions, first via line scanning^51–53^, and more recently through 2D imaging^48,50^, revealed fine structure in BOLD signals^54^. A recent ultrafast resting-state fMRI study highlighted the wealth of information embedded in spontaneous signals, revealing oscillations that drive long-range connectivity^49^ with neuronal origins^55^.

Here, we posit that the distinct spatiotemporal characteristics of FF and FB signals can be disentangled from ultrafast fMRI signals coupled. To model FF and FB signals, an effective bidirectional computational model is essential. We here propose that connective field (CF) modeling^56–59^ – unlike conventional functional connectivity (FC) – can capture directional specificity by leveraging CF size as a measure of spatial pooling, which varies across cortical layers. This unique bidirectionality allows CF modeling to resolve FF and FB information flow, even during resting-state (RS) activity, where both signals are more balanced than during stimulation. While prior CF applications have been limited to connectivity between cortical areas via FF processing and could not differentiate bottom-up from top-down sources^59–61^, we here introduced an ultrahigh spatiotemporal resolution fMRI approach combined with a layer-specific CF (lCF) model or characterizing the directionality of information flow between cortical layers (Figure 1). The ensuing lCF fMRI enables, for the first time, the prediction of neural activity in a specific voxel based on aggregate information from the layer it samples or to which it sends information. Non-invasive and network-wide in scope, the lCF model uniquely dissociates source and target layers, offering the ability to reveal specific FF and FB patterns noninvasively with fMRI, both in task-based and in RS activity.

**Figure 1.**
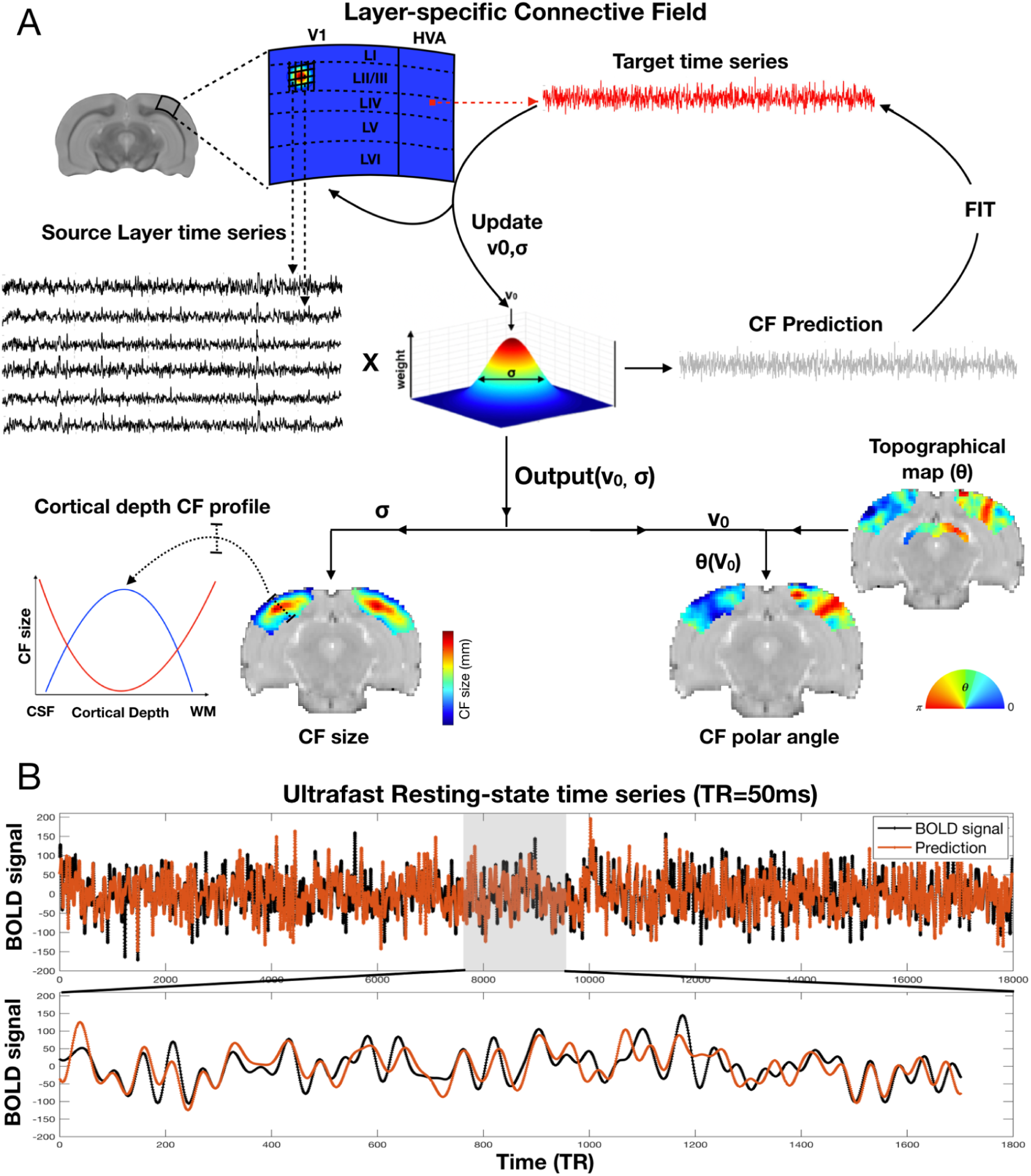
A: Layer CF Model. The lCF model predicts the neural activity of a recording site (voxel) in a target layer (e.g., layer IV of higher visual order areas (HVA)) based on the aggregate activity of a source layer or region (e.g. layer II/III of V1). Each voxel’s fMRI response is predicted using a 2D circular Gaussian CF model. The output parameters of the CF model are its center position (v_0_) in brain space and size (σ) within the source cortical layers. With a defined CF position and size, a predicted time-series is generated by weighting the CF model with the source BOLD time series. Optimal CF parameters are determined by minimizing the residual sum of squares between the predicted and actual time-series data. Our hypothesis was that, since lCF size reflects the extent of information integration, it would vary across cortical depth (layers). Specifically, we expected FF connections to exhibit an inverse U-shaped pattern, given that visual input is primarily processed in layer IV (LIV), while FB connections would follow a U-shaped pattern. **B:** Example trace of time series measured (black) and predicted (red) of a HVA voxel of layer IV based on aggregate activity from V1 LII/III.

## 2. Results

We developed the bidirectional lCF model to resolve FF and FB information, and apply it first to investigate information flow in the rat visual pathway. We begin by investigating area-to-layer projections from pre-cortical areas to V1, revealing the accuracy of the model in unravelling which layers receive FF signals from subcortical areas (Section 2.1). Next, we examined layer-to-layer communication between primary and higher cortical areas (Section 2.2), revealing unique signatures for both FF and FB signals across cortical areas. We then demonstrated that our findings in the visual system generalize to other cortical areas (Section 2.3) and finally, we applied our lCF approach to assess plasticity following network perturbations (Section 2.4).

### 2.1 Area-to-layer connective fields reveal the bottom-up flow of information in the geniculate and extrageniculate pathways

We first examine FF and FB signaling in the rodent visual system, which consists of two main pathways: the geniculate pathway, where V1 receives direct input from the lateral geniculate nucleus (LGN), primarily targeting LIV, and the extrageniculate pathway, where the lateral posterior nucleus (LP) projects to higher visual areas (HVA) in LIV and LV (Figure 2A). To assess whether fMRI can capture these pathways, we conducted multislice acquisitions of the visual system during RS and applied the lCF model to quantify projections from LGN and LP to different visual cortex layers. The CF model predicts activity in one region (e.g., V1 layers) based on the aggregate activity in another region (e.g., LGN), typically using a Gaussian kernel (Figure 1) to weight and integrate the time series, creating a predicted target signal. The center of the CF represents the location in brain space that has the highest correlation with the target area, while the size of the CF indicates the extent of information integration. CF models have been shown to capture topographically specific BOLD responses^59,60^. Figure 2C-D illustrates the CF polar maps obtained through the projection of retinotopic coordinates, measured in a separate scanning session where retinotopic mapping was performed, onto the CF locations (v_0_). The CF polar angle reveals a structured, retinotopic pattern across the entire visual pathway when the source is defined as either LGN or LP. The retinotopic pattern obtained from CF is very similar to the one obtained from retinotopic mapping (Figure S2). Notably, in the VC, the phase reversal points of the CF polar angle align with the borders between visual areas. This finding suggests that, even in the absence of direct visual input, functional connectivity preserves a visuotopic organization.

**Figure 2.**
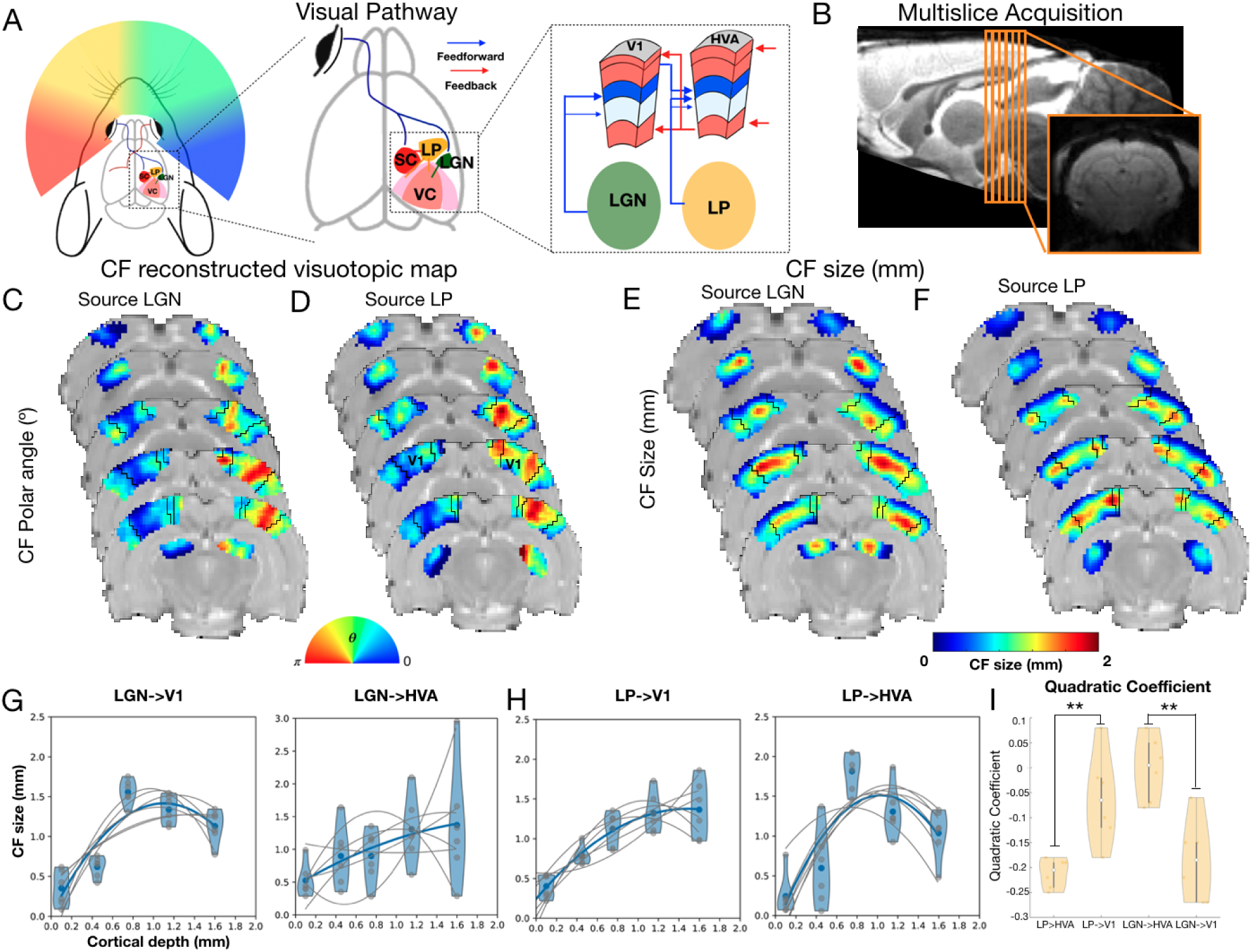
Area-to-layer CF profiles obtained for FF projections. **A**: Schematic illustration of the visual pathway, highlighting primary structures involved in FF and FB connections. **B:** Slices acquired in the multislice ultrafast RS fMRI datasets and representative functional slice. **C-F:** Visualizations of CF polar angle (C,D) and CF size (E,F) averaged across participants projected onto brain slices, with LGN (C,E) and LP (D,F) serving as the source areas. Black lines define the boundaries of VC areas. **G-H:** CF size profile (normalized to the mean) across the cortical layers of V1 and higher visual order areas (HVA), with LGN and LP serving as the source areas, respectively. The gray dots represent the data from each individual animal and gray lines the parabolic fit per animal. The blue dots and lines represent the median of all animals. **I:** Quadratic Coefficients obtained for the lCF size profile across cortical layers. The *** represents a *p*-value <0.001, ** *p*-value <0.01, and * *p*-value <0.05.

Figure 2E-F shows distinct projection patterns from the LGN and LP to different areas and layers of the VC, highlighting specialized pathways for visual information processing. When examining projections from the LGN to V1, we observed that the largest cortical CF sizes are concentrated in the middle layers, particularly layer IV, as shown in Figure 2E. This finding is further quantified in Figure 2G, which demonstrates an inverted U-shape profile of CF sizes, peaking in layer IV and decreasing in both the superficial and deeper layers. This characteristic profile suggests that LGN inputs are precisely targeted to the middle layers of V1. Notably, this inverted U-shape pattern is absent in the LGN projections to HVA, indicating a more specialized and layer-specific role for LGN inputs in V1. In contrast, LP projections show a complementary pattern. LP primarily feeds information to higher visual areas rather than V1. In V2, the LP projections exhibit an inverted U-shape CF profile, similar to the LGN to V1 pattern, peaking in the middle layers (Figure 2H). LP projections to V1 as well as LGN projections to V2 display a linear increase in CF size with cortical depth. A one-way ANOVA analysis of the quadratic coefficients of the functions describing CF size profiles across cortical depth revealed that the coefficients for LGN→V1 significantly differ from those for LGN→HVA (F(1,10)=18.87, p=0.0015), as well as LP→HVA differed from LP→V1 (F(1,10)=14.68, p=0.003) (Figure 2I). However, no significant differences were observed between LP→HVA and LGN→V1, nor between LP→V1 and LGN→HVA.

Furthermore, the quadratic coefficients obtained for LGN→V1 and LP→HVA were statistically different from 0 (LGN→V1 t(5)=5.6, p=0.003; LP→HVA t(5)=-17.7 p=1×10^−5^), while the coefficients obtained for LGN→HVA (t(5)=-0.06, p=0.95) and LP→V1 (t(5)=-1.7, p=0.16) were not significantly different from zero. Note that the CF size is independent from the population receptive field size. The pRF sizes are larger in HVAs (Figure S2), and in V1 pRFs tend to be larger in superficial and deeper layers (Fig. S2^47^).

Importantly, high temporal resolution also improves the accuracy of the CF estimation. Figure S3 demonstrates that the variance explained of the lCF model when applied to ultrafast data has an increase two to four fold compared with standard temporal resolution fMRI data.

Brain connectivity is typically analyzed using FC, which, unlike CF, does not account for spatial pooling between brain regions. As a result, FC has traditionally been considered unidirectional. However, FC strength could, in principle, help distinguish between FF and FB pathways based on characteristic FC strength profiles across cortical layers. For instance, if FC strength between LGN and V1 peaks in the middle layers of V1, this may indicate FF signaling. To determine whether the inverted-U-shaped, layer-specific CF profile, which reflects FF signaling, can be inferred from conventional FC methods, we computed the FC between the LGN, SC, LP, and V1 layers (Figure S1A and Table S1). The FC profile across cortical depth for all three connections (LGN, SC, and LP to V1 layers) did show an inverted-U trend, with the highest correlation coefficients observed in the middle V1 layers. However, the parabolic coefficients for these trends were very shallow (LGN→V1:-0.02; LP→V1:-0.01; SC→V1:-0.01), and none were statistically significantly different from zero (LGN→V1:F(5)=-2.13, p=0.08; LP→V1:F(5)=-0.67, p=0.53; SC→V1:F(5)=-1.16, p=0.3). Additionally, there were no significant differences in the parabolic coefficients among the LGN→V1, LP→V1, and SC→V1 pathways (F(2,17)= 0.86, p=0.44). In contrast, a comparison of the parabolic coefficients between the lCF size profile (Table S2) and FC profiles (Table S1) revealed a strong and highly significant difference (F(1,11)=21.85, p=0.0009). The parabolic curve associated with the lCF size profile was much more pronounced, with average parabolic coefficients of −0.19 for LGN→V1 and −0.21 for LP→HVA.

### 2.2 Layer-to-layer connective fields across cortical depth resolve FB and FF signals in spontaneous activity

Using single-slice acquisitions with a ultrahigh spatiotemporal resolution (120 µm x 120 µm) at 55 ms during spontaneous activity (where FB is enhanced and FB>FF^62^) and continuous binocular visual stimulation (FF enhanced, FF>FB^22^), we computed layer-to-layer CFs between V1 and V2 layers (Figure 3D-K). Layer-to-layer CF sizes revealed two distinct connectivity profiles across cortical layers that align with layer-specific FF and FB patterns: V1 projects information to the middle layers of V2 (inverted-U-shape profile), while the deep layers of V2 predominantly relay FB information to the superficial and deep layers of V1 (U-shaped profile, Figure 3G,K).

**Figure 3.**
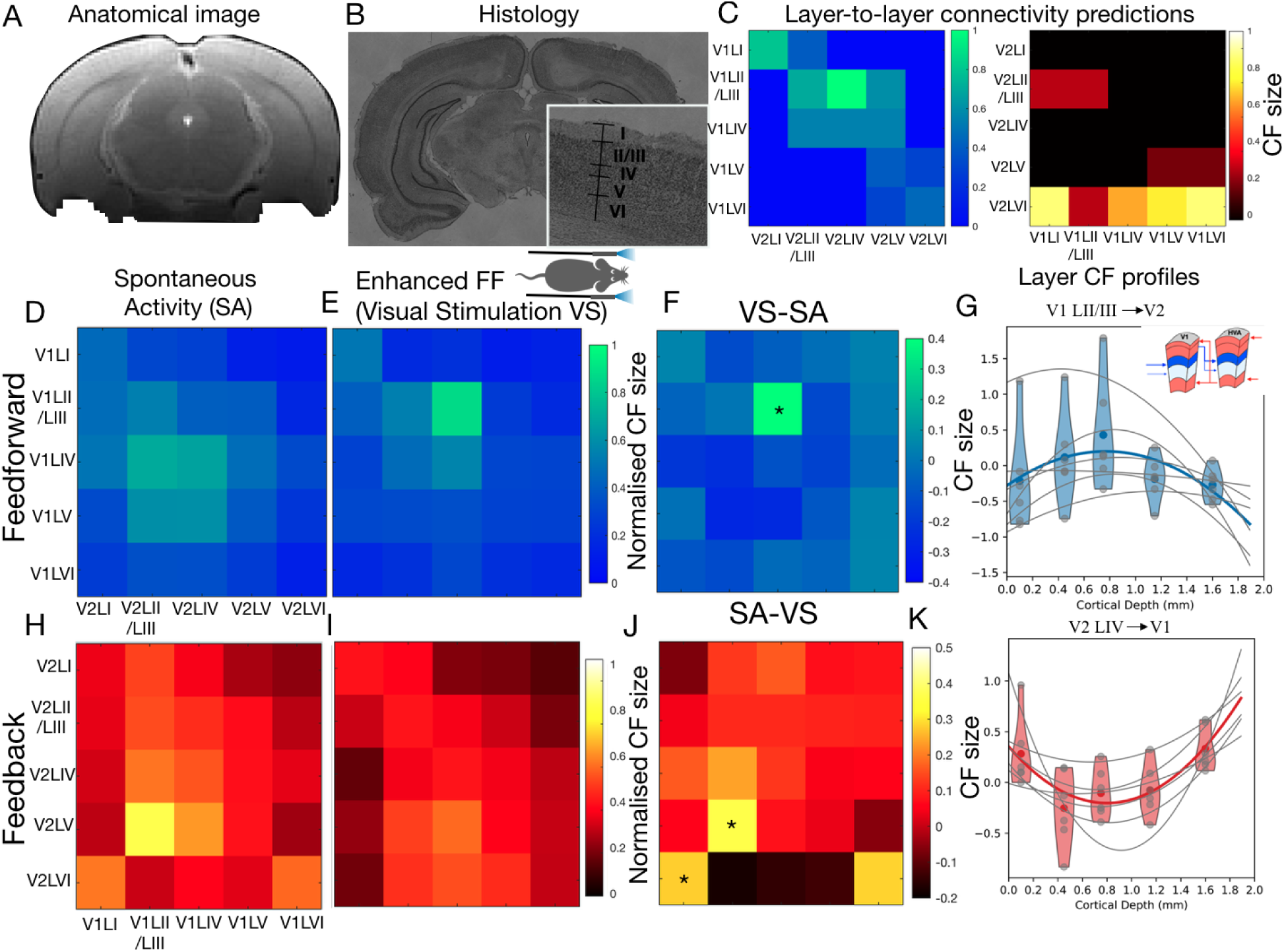
Layer-to-layer CF profiles. **A**: High-resolution anatomical image. **B:** A histological view delineating cortical layers. **C:** Schematic representation of the expected layer-to-layer connections based on the literature. FF connections (V1->V2) are indicated in the blue matrix and FB connections are shown in a red/yellow colorbar (V2->V1) (Federer et al 2021; ^63^**. D-F:** FF connectivity maps averaged across participants from V1 to V2 layers obtained under spontaneous activity (Panel D) and visual stimulation (Panel E), as well as the difference map subtracting spontaneous activity from visual stimulation non-normalised data, thereby isolating FF-specific pathways (Panel F). **G:** The CF size profile (normalized to the mean) illustrates how V1 layer II/III projects across all cortical layers in V2. The gray dots represent the data from each individual animal and gray lines the parabolic fit per animal. The blue dots and lines represent the median of all animals. **H-J:** FB connections averaged across participants between V2 to V1 layers acquired under spontaneous activity (Panel H) and visual stimulation (Panel I) conditions, with the visual stimulation data subtracted from RS data to isolate FB-specific pathways (Panel J). The subtraction was performed in the non-normalized maps. **K:** The CF size profile (normalized to the mean) demonstrates projections from V2 layer VI to all V1 cortical layers. The gray dots represent the data from each individual animal and gray lines the parabolic fit per animal. The red dots and lines represent the median of all animals. The * represent statistically significant differences between visual evoked and spontaneous activity.

In the context of FF projections from V1 to V2, we observed distinct lCF patterns between spontaneous and visual stimulation conditions. During spontaneous activity, the most prominent connections – large pooling extent (large lCF sizes)-were from V1 layer IV to V2 LII/III and LIV (Figure 3D). Under visual stimulation, the large lCF sizes were measured from V1 LII/LIII to V2 LII/LIII and LIV, and from V1 LIV to V2 LIV (Figure 3E). To differentiate the specific effects of FF and FB, we subtracted the spontaneous connectivity lCF size maps from those obtained during visual stimulation (Figure 3F). This subtraction revealed pronounced connections between V1 LI and V2 LI, as well as between V1 LII/LIII and V2 LIV. The lCF size obtained for the projection of V1 LII/LIII to V2 LIV was significantly larger during visual stimulation than during spontaneous activity (F(1,11)=8.92, p=0.0136). Quantifying these connections showed that the CF sizes from V1 LII/III to all V2 layers exhibit an inverse U-shaped pattern, with the largest sizes in V2 layer IV (Figure 3G).

For FB connections from V2 to V1, spontaneous activity data showed strong projections (larger CF sizes) from V2 LVI to V1 LI and LVI, and from V2 layer V to V1 LII/III and LIV (Figure 3H). By subtracting the visual stimulation lCF size maps from the spontaneous activity maps, we identified stronger sampling (larger CF sizes) between V2 layer VI and V1 LI (F(1,11)=6.99, p-value=0.02) and VI (F(1,11)=4.25, p-value=0.06), and between V2 layer V and V1 LII/III (F(1,11)=9.18, p-value=0.01) (Figure 3J). This was further quantified by analyzing the CF size profile across cortical depth from V2 LVI to all V1 layers, which exhibited a U-shaped profile, with CF sizes peaking in V1 LI and LVI (Figure 3K).

The layer-to-layer FF and FB connectivity maps, obtained with high spatiotemporal resolution, exhibit patterns similar to those derived from high temporal resolution (50 ms) and standard spatial resolution during spontaneous activity (Figure S4). Notably, these maps were acquired from different animals and scanning sessions, demonstrating the reproducibility of the lCF model. Additionally, Figure S4 presents FF and FB connectivity maps from higher-order visual areas both of which demonstrate comparable connectivity profiles: FF predominantly links middle layers, while FB primarily connects deeper layers.

Importantly, the layer-to-layer connectivity profiles obtained using lCF align with the projections identified in electrophysiology and calcium imaging studies using retrograde viral tracing and post-mortem quantification of axonal projections^63–65^, as shown in Figure 3C. Figure 3C shows a strong FF projection between V1 LII/LIII and V2 LIV; and strong FB projection between V2 LVI and V1 LI and LIV.

FC strength is typically calculated using Pearson correlations, meaning that layer-to-layer FC between V1 and V2 is symmetrical (i.e., FC(V1→V2) is indistinguishable from FC(V2→V1)). As a result, this approach cannot disentangle the direction of information flow. Figure S1B illustrates the layer-specific FC calculated between V1 and V2 layers, revealing FC map V2→V1 is symmetrical to FC V2→V1 and that FC profiles differ substantially from those obtained using the lCF model both for the FF connections (V1LV->V2LIV, V1LV->V2LV, V1LVI->V2LIV, V1LVI->V2LV) as well as for FB connections (V2LII/III->V1LVI, V2LIV->LV, V2LIV->LVI, V2LV->V1LII/III, V2LV->V1LV, V2LV->V1LVI; V2LVI->VILI, V2LVI->VILVI).

### 2.3 Layer CF reveals directional connectivity in the somatosensory and motor cortices

To investigate whether the CF size profiles for FF and FB pathways observed in the visual system are general (i.e., they exhibit similar properties in other brain regions) we analyzed area-to-layer and layer-to-layer CFs in the somatosensory and motor pathways from multislice and single-slice ultrafast data, respectively. Given that the cortex in somatosensory and motor areas is thicker than in the VC, we were able to define cortical layers with greater precision using standard spatial resolution data (Figure 4B).

**Figure 4.**
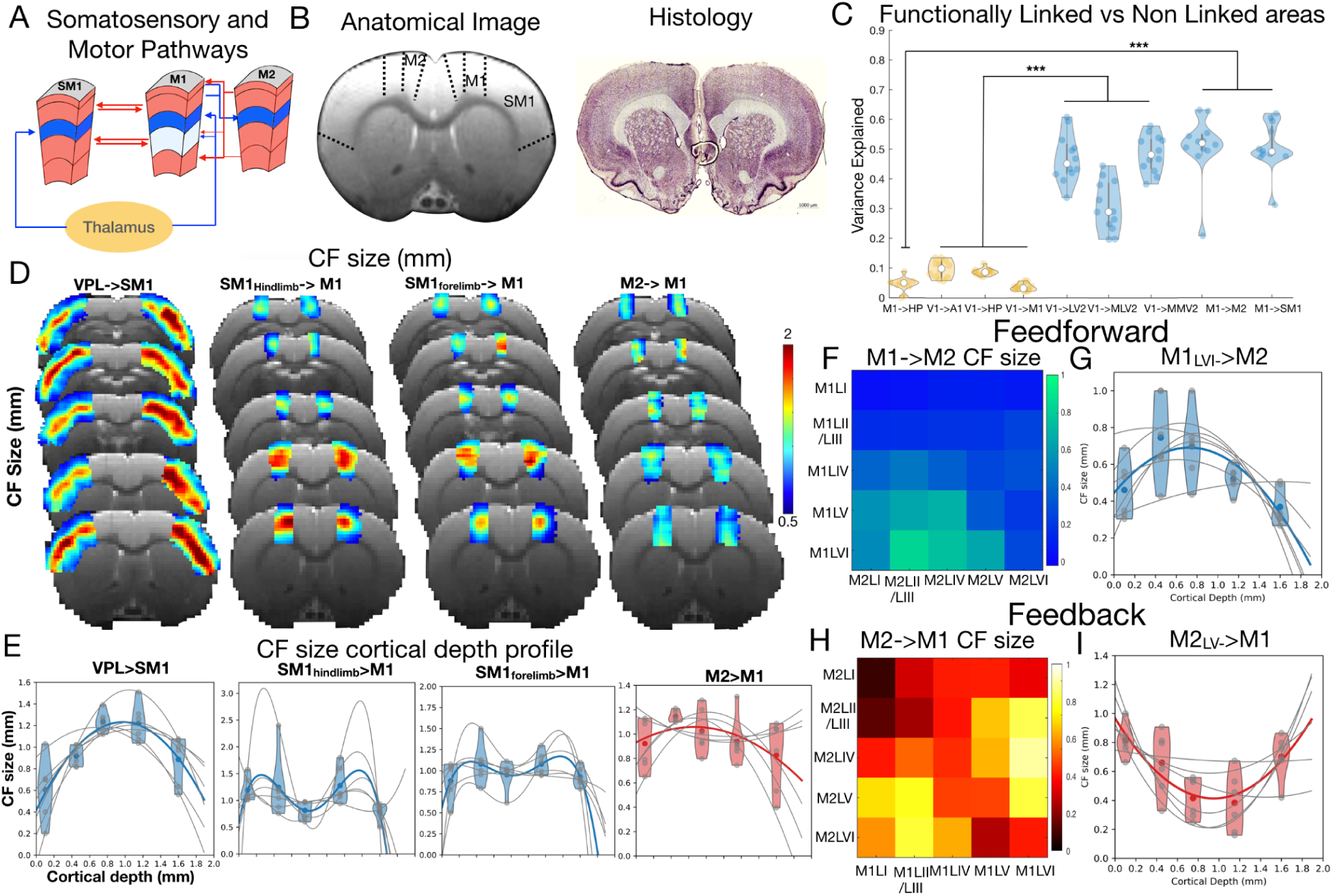
Connectivity in the somatosensory and motor cortices. **A**: Schematic representation of the somatosensory and motor pathways. **B:** Anatomical image and histological slice of the somatosensory and motor cortices. **C:** Variance explained obtained from brain regions anatomically and/or functionally connected (blue) and brain regions that are not directly connected (yellow) (A1: Primary Auditory Cortex; HP: Hippocampus). The *** represents a *p*-value <0.001, ** *p*-value <0.01, and * *p*-value <0.05. **D:** Visualizations of CF size averaged across participants projected into the brain slice for the following connections: VPL->SM1; SM1(forelimb and handlimb)->M1; and M2->M1. **E:** CF size profile (normalized to the mean) across the cortical layers of SM1 and M1 for the following connections: VPL->SM1; SM1(forelimb and handlimb)->M1; and M2->M1. The gray dots represent the data from each individual animal and gray lines the parabolic fit per animal. The blue or red dots and lines represent the median of all animals. **F:** FF connections between M1 and M2 layers acquired under spontaneous activity. **G:** The CF size profile (normalized to the mean) demonstrates projections from M1 layer VI to all M2 cortical layers. **H:** FB connections between M2 and M1 layers acquired under spontaneous activity. **I:** The CF size profile (normalized to the mean) demonstrates projections from M2 layer V to all M1 cortical layers.

In both the primary somatosensory cortex (SM1) and primary motor cortex (M1), layer IV receives direct input from thalamic structures, such as the ventral posterolateral nucleus (VPL) (Figure 4A). When VPL was set as the source, CF size profiles showed larger values in layer IV of SM1, with an inverse U-shaped profile that peaks at this layer (Figure 4D-E), consistent with findings in the visual pathway (Figure 2E-H). Additionally, SM1 and M1 exhibit strong connectivity through LII/III and LV (Figure 4A). When examining the CF size profiles across these layers, we observed slightly larger CF size in LII/III and LV than in LI, LIV and LVI (Figure 4D,E). The CF size profile between the secondary motor cortex (M2) and M1, despite mostly carrying FB signals, also displayed an inverted-U-shaped profile on average, although with a shallow curvature and high variability between animals.

The layer-to-layer connections between M1 and M2 (Figure 4F) mirror those observed in the visual pathway, with deep layers of M1 projecting predominantly to the middle layers of M2. Quantification of CF size from M1 LVI to all M2 layers reveals a ∩-shaped profile, with CF size peaking in LII/III and IV of M2 (Figure 4G). Conversely, the connections from M2 to M1 show the opposite pattern: deep layers of M2 feed information to the superficial layers of M1, while middle layers of M2 (LII/III, LIV, and LV) project to the deep layers of M1 (Figure 4H). The CF size profile across cortical depth, with M1 layer V (LV) feeding into all M1 layers (Figure 4I), further emphasizes this layer-specific communication pattern.

The clarity of the layer-to-layer connectivity profiles improves as the temporal resolution of data acquisition increases (Figure S5). For both the VC and motor cortex slices, we downsampled the temporal resolution from 50 ms to 350 ms. Results revealed that as temporal resolution decreases, the distinction between FF and FB connectivity patterns becomes less pronounced. At 350 ms, FF connections are no longer distinguishable from FB, with FF patterns dominating the data (Figure S5). The loss of specificity in layer-to-layer connectivity maps with decreased temporal resolution is evident in both visual and somatosensory/motor systems (Figure S5). This lack of precision may explain why the area-to-layer projections from M2 to M1 (obtained from multislice data, where TR = 350 ms), on average, do not exhibit a U-shaped profile (Figure 4E). These findings underscore the critical importance of high temporal resolution for accurately capturing the nuanced dynamics of cortical connectivity.

Furthermore, to validate the biological relevance of the CFs, we computed them between brain regions with known anatomical connections, specifically between visual areas and between motor and somatosensory areas, as well as between regions where strong connections are not expected, such as between the VC and auditory/motor cortex, or the VC and hippocampus. Brain regions with known anatomical connections have much higher variance explained than regions that are not anatomically or functionally connected (Figure 4C). This difference in variance explained is highly significant (F(1,70)=326.21, p<0.001).

### 2.4 Layer CF reveals circuit reorganization following cortical lesions

The ability of lCF estimates to distinguish between FF and FB pathways opens the possibility for mapping plasticity upon network perturbations, reinforcing its causal validity rather than merely reflecting correlational effects. To test this, we induced cortical blindness (CB) by bilaterally lesioning the V1 region in seven animals (Figure 5A). Figure 5B,C displays anatomical and histological images of the lesions, with V1 appearing lighter and whitish in color, indicating cell death. The layered structure of the cortex is also less clear. These lesions disrupt the visual pathway and we expected these changes to be reflected in the CF size profiles. Specifically, we hypothesized that in the absence of an intact visual pathway, the LGN—which normally projects directly to V1—would instead project to HVA. Additionally, we anticipated that the extrageniculate pathway (LP→HVA) would strengthen its projections, compensating for the loss of the geniculate pathway.

**Figure 5.**
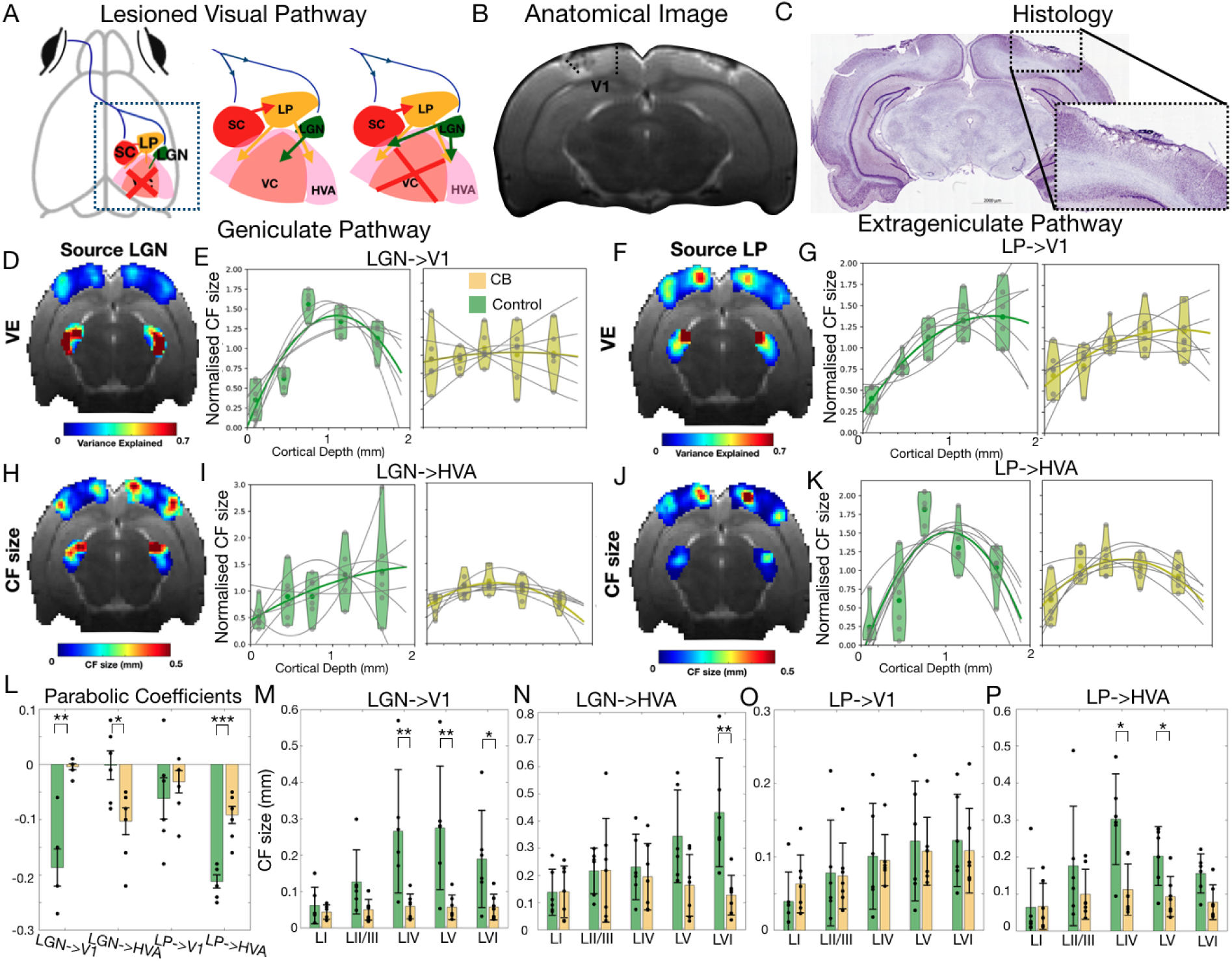
Lesions. **A**: Schematic representation of the V1 lesion and expected reorganization of the pathway. **B:** Anatomical image illustrating the bilateral V1 lesions. **C:** Histological analysis of a V1 lesioned animal brain. **D** and **F:** Variance explained of the CF model averaged across CB animals and computed taking as source LGN and LP, respectively, projected on brain slices. **H** and **J:** CF size averaged across CB animals when LGN and LP are taken as sources, respectively, projected on brain slices. **E, G, I, K:** Quantitative analysis of the CF size (normalized to the mean) as a function of cortical depth for HC (green) and for CB animals (yellow) obtained for the connections: LGN->V1 (Panel E); LP->V1 (Panel G); LGN->HVA (Panel I); LP->HVA (Panel K). The gray dots represent the data from each individual animal and gray lines the parabolic fit per animal. The green or yellow dots and lines represent the median of all animals from each group. **L:** Quadratic Coefficients obtained for the lCF size profile across cortical layers for controls (green) and CB animals (yellow). The *** represents a *p*-value <0.001, ** *p*-value <0.01, and * *p*-value <0.05. **M, N, O, P:** Average CF size per layer calculated for HC and CB animals obtained for the connections: LGN->V1 (Panel M); LP->V1 (Panel N); LGN->HVA (Panel O); LP->HVA (Panel P). The non-normalized lCF sizes are shown in Figure S6.

We calculated the area-to-layer CFs seeding from the LP and LGN to predict the VC time series (Figures 5D-P). The variance explained of the lCF projecting from LGN and LP to V1 in CB animals was close to zero (VE: LGN→V1=0.04± 0.02; LP→V1=0.03±0.02, Table S3) and highly reduced compared to healthy controls (HC) animals (VE: LGN→V1=0.42±0.05; LP→V1=0.22±0.07) confirming that CFs capture biologically relevant information (Figure 5D,F,H,J). We then analyzed CF size profiles across cortical depth. Meaningful FF projections like LP to V2 (in both HC and CB) and LGN to V1 (in HC) displayed an inverted-U-shaped profile. Notably, for the LGN to V1 connection in CB animals, this profile was absent due to the lack of neural activity in V1 (Figure 5E). Therefore, the parabolic coefficients from the fits of the CF size across cortical layers are highly significantly different between HC and CB animals (F(1,11)=28.92, p=0.0003), Figure 5H. In addition, there are significant changes in CF size in V1 LIV (F(1,11)=8.63, p=0.01), LV (F(1,11)=9.84, p=0.01) and LVI (F(1,11)=5.81, p=0.04) between HC and CB animals when the source is LGN (Figure 5M).

Interestingly, the LGN to HVA connection in CB animals showed an emerging an inverted U-shaped pattern (Figure 5I, N), which was not present in HC (linear trend), suggesting potential compensatory changes in the visual pathway following the V1 lesion. This resulted in significant changes in the parabolic coefficient from the fits of the CF size across cortical layers between HC and CB animals (F(1,11)=4.87, p=0.05, Figure 5L), with CB animals showing a more negative parabolic coefficient. Furthermore, there were significant changes in CF size in HVA layer IV between HC and CB animals when the source is LGN (F(1,11)=11.65, p=0.007) (Figure 5N).

In contrast, non-meaningful connections, such as LP to V1, exhibited a linear increase in CF size with cortical depth for both HC and CB groups, with no significant differences in the parabolic coefficients (F(1,11)=0.04, p-value=0.85), neither on the average CF size obtained per layers (p-value>0.05) between HC and CB animals (Figure 5G,L,O).

The projection from LP to V2 shows an inverted-U-shaped profile for both HC and CB, however, the curvature of the parabolic fit obtained for the CF is shallower than the one obtained for the HC (F(1,11)=30.3, p=0.003, Figure 5L). The CF size in layer IV and V is significantly higher in HC than in CB animals (F(1,11)=8.55, p=0.015; F(1,11)=7.33, p=0.022, respectively, Figure 5P).

Two additional animals were monocularly lesioned. Figure S7 shows that in the spared hemisphere the lCF size calculated from LGN to V1 shows larger CF sizes in the Layer IV. In the lesioned site, CF size and VE are significantly decreased (VE∼0) and the CF size increases in the HVAs surrounding the lesion.

## 3. Discussion

We introduced Layer-resolved connective fields derived from ultrahigh spatiotemporal resolution fMRI, and showed that their connectivity signatures are associated with FF and FB processes across cortical layers. To our knowledge, this is the first time that FF and FB signals were dissociated from spontaneous activity non-invasively with fMRI, thereby opening new ways of characterising information flow in the brain in general. As predicted in^56^, we found that CF size – a measure of information integration – produces two distinct connectivity profiles across cortical layers. FF connectivity was characterized by a ∩-shaped pattern, with larger CF sizes concentrated in LIV; while FB connectivity exhibited a U-shaped profile, with CF sizes peaking in the superficial (LI) and deep (LVI) cortical layers. The lCF size profiles are in line with our understanding of underlying neural functional and anatomical architecture, i.e. axonal projections^40,66^. Importantly, we have shown that lCF profiles associated with FF and FB are general, as observed in different brain sensory systems (somatosensory, motor and visual), and that they provide meaningful insight into information flow upon injury (as observed in the CB animal model), suggesting it may have high relevance for understanding disease progression, recovery from injury, development, and plasticity, among other applications.

FF signals are generally considered to carry sensory information, while FB signals reflect top-down expectations or predictions^67,68^. However, the ability of lCF to dissociate FF and FB signals during spontaneous activity challenges the notion that FF signals are exclusively stimulus-dependent. Instead, the dissociation of FF and FB signals during spontaneous activity suggests that laminar-specific FF and FB interactions are continuously active and interacting. This is consistent with studies showing that cortical responses to natural stimulation closely resemble those observed during spontaneous activity^62^ and that the network structure distinguishing FF and FB-specific neurons persists even without stimulation ^69^. These findings support the view that spontaneous activity reflects the underlying connectivity and the balance between FF and FB signals.

The area-to-layer CF patterns primarily reflect the FF pathway, highlighting the complementary roles of the geniculate (retina→LGN→V1) and extrageniculate (retina→SC→LP→HVA) pathways in visual processing^70^. The LGN and LP provide targeted input to the middle layers of V1 and higher visual areas (HVA), respectively. Additionally, CF maps derived from spontaneous activity in both geniculate and extrageniculate pathways demonstrated a clear visuotopic organization. This underscores that CF is a stimulus-agnostic tool, allowing for the study of sensory topography and the underlying bottom-up and top-down mechanisms in the absence of external stimulation^56,71^. This finding is consistent with CF studies in humans, where similar retinotopic maps were estimated from both task-based data (e.g., movie watching) and spontaneous activity data^59,60^.

Furthermore, for animals with bilateral cortical lesions in V1 – CB, the lCF estimates revealed that HVAs direct FF input from the LGN bypasses V1 and follows an inverted U-shape CF size profile. This finding aligns with invasive V1 lesion studies in both rodents and humans. In rodents, it has been shown that after a V1-induced lesion, LGN neurons project directly to several HVAs^70,72^. Similarly, in humans, residual vision — such as blindsight — following V1 injury is thought to be mediated by circuits that bypass V1, directly relaying signals to higher-order cortical areas^73^. These results suggest that the distinct properties of these visual areas depend on inputs from regions beyond V1, indicating the existence of an adaptive alternative pathway that maintains visual processing despite the loss of primary visual input.

Our work highlights the essential role of ultrahigh temporal resolution for disentangling FF and FB patterns as well as generally bidirectional computational models of information flow within cortical networks. This finding is in line with previous studies that have shown that increasing fMRI temporal resolution improves the performance of directional sensitive models, i.e., Granger Causality^74^. This suggests a paradigm shift in the current fMRI studies attempting to dissociate FF and FB signals, which currently focus on layer selectivity (high spatial resolution) overlooking the temporal resolution. However, we acknowledge that sacrifices of spatial coverage may be necessary to achieve this temporal resolution, although parallel mapping methods^75–79^ are likely to become highly useful for achieving both spatial and temporal high resolution, simultaneously.

This study has three main limitations. First, isolating FB signals, i.e. using attentional or imagery tasks, is challenging in rodents and preclinical scanners. While FF and FB signals coexist during both stimulation and spontaneous activity, an FB-specific task could help confirm the U-shaped FB profile. However, previous research suggests spontaneous activity is predominantly FB-driven^62^. Second, sedation likely weakens neural activity in higher-order areas, potentially suppressing FB signals and biasing lCF estimates. Although the animals were sedated throughout, a study by Semedo et al. found FB responses persist under anesthesia, suggesting FB activity is not entirely abolished^22^. Third, V1 lesions may have affected other cortical areas and the vascular network, potentially altering BOLD signals in ways mistaken for plasticity. Future studies could refine FB investigation by optogenetically stimulating V2, isolating FB mechanisms without influencing FF activity.

Non-invasively disentangling FF and FB signals in rodents enables pathway-wide investigations without requiring viral injections or electrode placements, which may alter neural activity. With advancements in high-resolution MRI, this approach could extend to high-field clinical scanners. The ability of lCF to non-invasively dissociate FF and FB connections from spontaneous activity offers exciting opportunities to explore the FF/FB balance in humans. Understanding the FF/FB dynamics is crucial, as disruptions contribute to conditions like schizophrenia, ASD, depression, ADHD, and PTSD. Tracking FF and FB signals could also enhance our understanding of action planning, attention, predictive coding, and contextual modulation. Clinically, detecting FF/FB imbalances may improve diagnostics and inform targeted interventions, such as neuromodulation or cognitive therapies, to restore proper communication.

## 4. Methods

All the experiments strictly adhered to the ethical and experimental procedures in agreement with Directive 2010/63 of the European Parliament and of the Council, and were preapproved by the competent institutional (Champalimaud Animal Welfare Body) and national (Direcção Geral de Alimentação e Veterinária, DGAV) authorities.

In this study, we conducted two distinct sets of experiments to investigate FF and FB circuits: a) in the visual cortex of healthy and cortically lesioned animals; and b) in the somatosensory and motor cortices of healthy animals.

Regarding the experiments probing the visual pathway, a total of 19 Long Evans rats (12 controls: 10 F, 12-30 weeks old, w=412±75 g; and 7 rats for the lesion experiment: 5 F, 8-24 weeks old, w=364±45 g) were scanned in two separate sessions. Session 1 focused on retinotopic mapping, while Session 2 involved ultrafast acquisitions. In Session 2, three sets of ultrafast experiments were conducted to investigate FF and FB connectivity in the visual system of rats: set 1 – Standard spatial resolution RS scans in control animals (N=6); set 2 – High spatial resolution scans (both RS and visual stimulation) in control animals (N=6); set 3 – Standard spatial resolution RS scans in the rats with lesions in the V1. Except for the cortical lesions experiment (set 3), Session 1 took place one week before Session 2.

In the experiments probing the somatosensory and motor cortices, six Long Evans Rats (4 F, 11-15 weeks old, w=423±69 g) were scanned.

### 4.1 Animal preparation

All in-vivo experiments were performed under sedation. The animals were induced into deep anesthesia in a custom box with a flow of 5% isoflurane (Vetflurane, Virbac, France) mixed with oxygen-enriched (27%) medical air for ∼2 min. Once sedated, the animals were moved to a custom MRI animal bed (Bruker Biospin, Karlsruhe, Germany) and maintained under ∼2.5–3.5% isoflurane while being prepared for imaging. Eye drops (Bepanthen, Bayer, Leverkusen, Germany) were applied to prevent the eyes from drying during anesthesia. Approximately 5 min after the isoflurane induction, a bolus (0.05 mg/kg) of medetomidine (Dormilan, Vetpharma Animal Health, Spain) consisting of a 1 mg/ml solution diluted 1:10 in saline was administered subcutaneously. Ten to eighteen minutes after the bolus, a constant infusion of 0.1 mg/kg/h of medetomidine, delivered via a syringe pump (GenieTouch, Kent Scientific, Torrington, Connecticut, USA), was started. During the period between the bolus and the beginning of the constant infusion, isoflurane was progressively reduced until 0%.

Temperature and respiration rate were continuously monitored via a rectal optic fiber temperature probe and a respiration sensor (Model 1025, SAM-PC monitor, SA Instruments Inc., USA), respectively, and remained constant throughout the experiment. Each MRI session lasted between 2h30min and 3h. At the end of each MRI session, to revert the sedation, 2.0 mg/kg of atipamezole (5 mg/ml solution diluted 1:10 in saline) (Antisedan, Vetpharma Animal Health, Spain) was injected subcutaneously at the same volume of the initial bolus.

### 4.2 MRI acquisition

All the MRI scans were performed using a 9.4T Bruker BioSpin MRI scanner (Bruker, Karlsruhe, Germany) operating at a ^1^H frequency of 400.13 MHz and equipped with an AVANCE III HD console and a gradient system capable of producing up to 660 mT/m isotropically. An 86 mm volume quadrature resonator was used for transmittance. A 20 mm loop surface coil and a 4-element array cryoprobe (Bruker, Fallanden, Switzerland) were used for signal reception during Session 1 and Session 2 of visual system experiments, respectively. The cryogenic reception coil was also used for the somatosensory and motor cortex experiments. The software running on this scanner was ParaVision^®^ 6.0.1.

After placing the animal in the scanner bed, localizer scans were performed to ensure that the animal was correctly positioned and routine adjustments were performed. B_0_ maps were acquired. A high-definition anatomical T_2_-weighted Rapid Acquisition with Refocused Echoes (RARE) sequence (TE/TR = 13.3/2000 ms, RARE factor = 5, FOV = 20 × 16 mm^2^, in-plane resolution = 80 × 80 μm^2^, slice thickness = 500 μm, t_acq_ = 1 min 18 s) was acquired for accurate referencing. Importantly, the functional MRI scans were started ∼30 min after the isoflurane was removed from the breathing air to avoid the potentially confounding effects of isoflurane ^80^.

### 4.3 Visual system experiments

#### 4.3.1 Session 1: Retinotopy

##### Acquisition

Functional scans were acquired using a gradient-echo echo-planar imaging (GE-EPI) sequence (TE/TR = 12/1500 ms, FOV = 20.5 × 20.5 mm^2^, resolution = 227 × 227 μm^2^, slice thickness = 788 μm, 8 slices covering the visual pathway, flip angle = 15°). The animals underwent a total of 6 runs of the retinotopic stimulus, each run taking 7 min and 39 s to acquire (306 repetitions). In addition, a 10 min RS scan was also acquired with the same parameters.

##### Stimulus and Setup

The visual stimuli consisted of a luminance contrast-inverting checkerboard drifting bar ^47^. The bar aperture was composed by alternating rows of high-contrast luminance checks moved in 8 different directions (4 bar orientations: horizontal, vertical and the two diagonal orientations, with two opposite drift directions for each orientation). The bar moved across the screen in 16 equally spaced steps, each lasting 1 TR. The bar contrast, width, and spatial frequency were 50%, ∼14.5°, and ∼0.2 cycles per degree of visual angle (cpd), respectively. The retinotopic stimulus consisted of four stimulation blocks. At each stimulation block, the bar moved across the entire screen during 24 s (swiping the visual field in the horizontal or vertical directions) and across half of the screen for 12 s (swiping half of the visual field diagonally), followed by a blank full screen stimulus at mean luminance for 45 s. A single retinotopic mapping run consisted of 246 functional images (60 pre-scan images were deliberately planned to be discarded due to coil heating).

The complex visual stimuli necessary for retinotopic mapping were generated outside the scanner and back-projected with an Epson EH-TW7000 projector onto a semitransparent screen positioned 2.5 cm from the animals eyes. The setup has been previously described ^47^. Visual stimuli were created using MATLAB (Mathworks, Natick, MA, USA) and the Psychtoolbox. An Arduino MEGA260 receiving triggers from the MRI scanner was used to control stimuli timings.

#### 4.3.2 Session 2: Standard spatial resolution RS in control animals

Three ultrafast RS datasets were obtained using GE-EPI acquisitions from: a multislice set; a VC slice and a visual pathway (VP) slice (Figure 6C).

**Figure 6.**
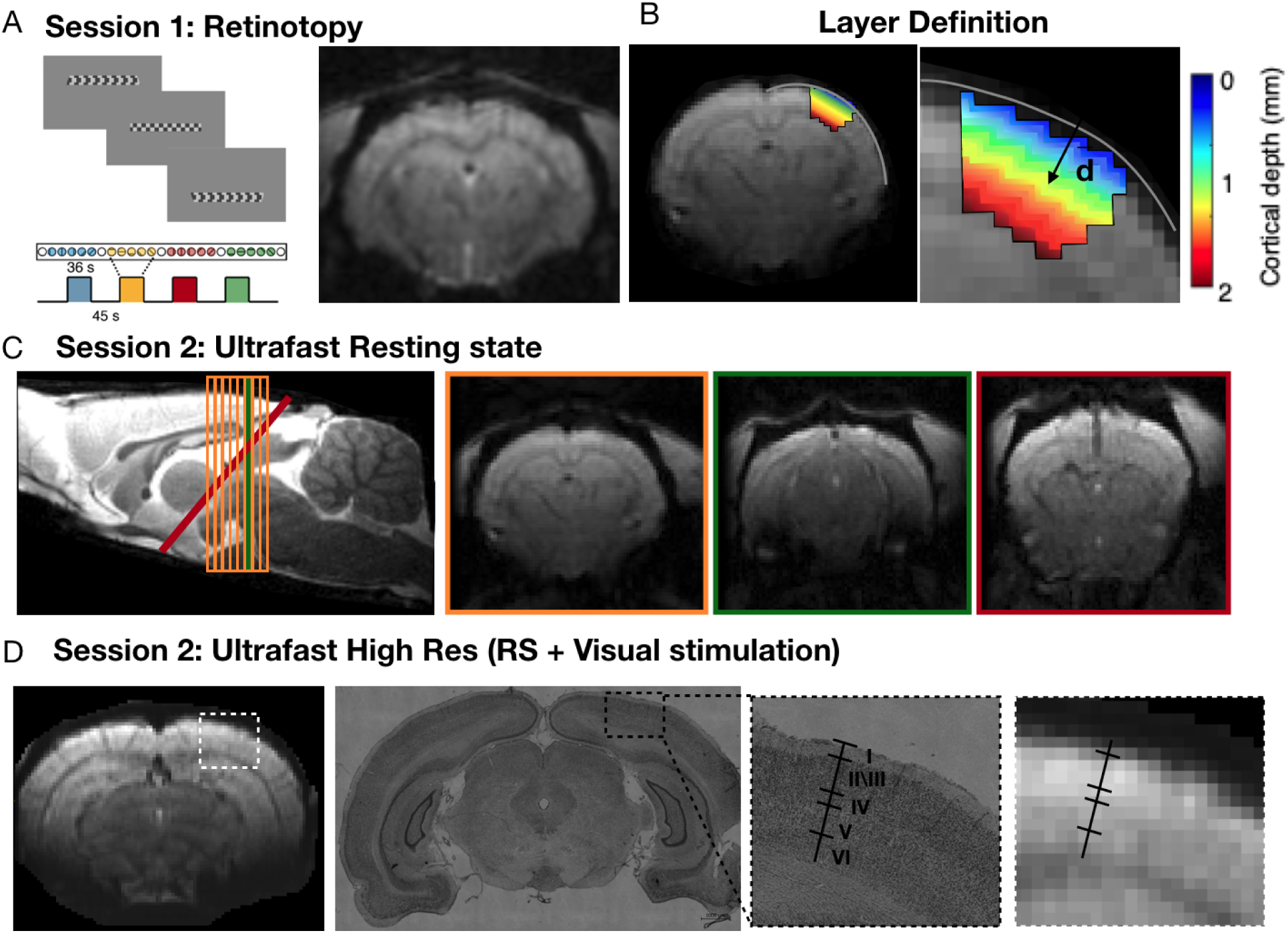
MRI acquisition protocol. **A**: Retinotopic stimulus, stimulation paradigm and an example of a slice acquired in Session 1. **B:** Representative functional slice acquired in Session 2 (Standard Resolution) and its respective layer definition (where d is the distance from cortical surface). **C:** Representative functional slices of the RS datasets (multislice (orange), single VC slice (green) and single VP slice (red)) acquired at standard resolution. **D:** High spatial resolution VC slice acquisition and histology analysis of a HC animal with the layer definition.

##### Multislice set

The functional MR imaging was acquired using a GE-EPI sequence (TE/TR = 12/350 ms, FOV = 20.5 × 20.5 mm^2^, resolution = 227 × 227 μm^2^, slice thickness = 788 μm, 8 slices, t_acq_ = 20 min 500 ms).

##### VC and VP slices

The functional MR imaging was acquired using a GE-EPI sequence (TE/TR = 11.6/50 ms, FOV = 20.5 × 20.5 mm^2^, resolution = 227 × 227 μm^2^, slice thickness = 788 μm, t_acq_ = 20 min 500 ms).

#### 4.3.3 Session 2: Ultra-high spatial resolution spontaneous activity and visual stimulation in control animals

Two high spatial resolution ultrafast datasets of the VC were obtained: one during RS and one during visual stimulation. The functional MR imaging was acquired from the VC slice defined above using a GE-EPI sequence (TE/TR = 11.6/55 ms, FOV = 20.5 × 20.5 mm^2^, resolution = 163 × 163 μm^2^, slice thickness = 700 μm, t_acq_ = 20 min 500 ms). To avoid contamination of the visual stimulation effect to the RS, the RS scan was acquired before the visual stimulation scan.

##### Visual stimulation

A bifurcated optic fibre connected to a blue LED (*λ* = 470 nm and I = 8.1 × 10−1 W/m^2^) was placed horizontally in front of each eye of the animal (spaced up to 1 cm) for binocular visual stimulation. The blue LED was connected to an Arduino MEGA260 receiving triggers from the MRI scanner and was used to generate square pulses of light. A stimulation frequency of 8 Hz was used. The LEDs flickered continuously during the entire duration of the scan.

#### 4.3.4 Session 2: Standard spatial resolution RS in lesioned animals

The MRI acquisition of lesioned animals was identical to the one obtained for controls (see Section 4.3.2). Session 1 took place 1 week prior to the lesion surgery and the animals were scanned 1 week post-lesion for Session 2.

##### V1 Lesions: Ibotenic acid

To investigate how FF and FB pathways were reorganized following V1 lesions, *n* = 7 animals were injected bilaterally with an ibotenic acid (Abcam, Romania) solution (excitotoxic agent, 1 mg/100 μL) in the V1. The acid was injected using a Nanojet II (Drummond Scientific Company). Animals were anesthetized with isoflurane (anaesthesia induced at 5% concentration and maintenance below 3%), and a scalpel incision was made along the midline of the skull, the skin retracted and the soft tissue cleaned from the skull with a blunt tool.

Coordinates for the craniotomies and amounts required for the injections were determined for each individual animal based on T2-weighted anatomical images acquired before the surgery. We injected in 5 different AP coordinates; two injection pulses on the one injection site for the first AP coordinate; and 4 injection pulses for each of the following injection sites, two per AP coordinate, for full coverage of V1. Each injection pulse was administered at a rate of 23 nL/s with 2–3 s between pulses, and each pulse consisted of 32 nL. Waiting time before removing the injection pipette after the last pulse was 10 min. The craniotomies were then covered with Kwik-Cast™ (World Precision Instruments, USA) and the scalp was sutured. Before the animals recovered from surgery, they were injected, subcutaneously, with 5 mg/Kg body weight of carprofen (Rimadyl ®, Zoetis, U.S.A). Following surgery, the animals were allowed to recover for around 7 days to avoid MRI acquisition artefacts due to potential inflammation and mechanical damage from the pipette.

#### 4.3.5 Histology

Histological analysis was performed for one control and two V1 lesioned animals. Animals were perfused transcardially, ∼24 h hours after Session 2. First with a PBS 1X solution, followed by 4% PFA. The brain was extracted and kept in 4% PFA for approximately 12 h. After this, the brain was placed in a 30% sucrose solution for a minimum of 4 days, after which the tissue was embedded in a frozen section compound (FSC 22, Leica Biosystems, Nussloch, Germany) and sliced on a cryostat (Leica CM3050S, Leica Biosystems, Nussloch, Germany). After sectioning, the brain slices were stained with cresyl violet, mounted with mowiol mounting medium, and imaged using a Zeiss AxioImager M2.

### 4.4 Somatosensory and motor system experiments

Regarding the experiments probing the somatosensory and motor pathways, a multislice set and a single slice covering the somatosensory and motor pathways were acquired during RS. The scanning parameters are the same as in the multislice and single slice ultrafast acquisitions described in Section 4.3.2.

### 4.5 Data Analysis

#### 4.5.1 Preprocessing

##### 4.5.1.1 Retinotopy data

Images were extracted from the scanner using the vendor image reconstruction pipeline and converted to Nifti. Outlier correction was performed (time points whose signal intensity was 3 times higher or lower than the standard deviation of the entire time course were replaced by the mean of the 3 antecedent and 3 subsequent time points).

##### 4.5.1.2 Ultrafast data

The complex data from each cryoprobe channel was extracted and transformed into k-space, where zero filling was removed. Afterward, the images were reconstructed back into image space, and outlier correction was carried out by manually identifying time points with average brain signals deviating ± 2–3 standard deviations from a 2nd-order polynomial trend. Subsequently, new voxel values for these outlier time points were estimated using piecewise cubic interpolation based on the remaining time points. This correction was performed slice by slice. Next, NORDIC denoising was applied with a sliding window of [5, 5, 1], treating each channel separately. After denoising, zero filling was reintroduced and the data from the four channels were combined to form the final magnitude images.

##### 4.5.1.3 Preprocessing steps common to the retinotopy and ultrafast data

All the following preprocessing steps were performed using a home-written Python pipeline using Nypipe.

Simultaneous motion and slice timing correction were performed using Nipy’s SpaceTimeRealign function ^81^. Brain extraction was done using AFNI function Automask applied to a bias field corrected (ANTs) mean functional image. The skull stripped images were inspected and upon visual inspection a further mask was manually drawn when needed using a home-written script. The skull stripped images then underwent co-registration and normalization to an atlas ^82^. Manual co-registration was performed using ITK SNAP, aligning the mean functional image of each run to the anatomical image. Normalization was performed by calculating the affine transform matrix that aligns each anatomical image with the atlas template using ANTs. The two sets of transform matrices were then refined and applied to all the functional images using ANTs.

Following normalization, for Session 1 data (retinotopy), the voxels’ signals were detrended using a polynomial function of factor 2 fitted to the resting periods and spatially smoothed using a 2D Gaussian Kernel (FWHM = 0.15 mm^2^). For Session 2 data and somatosensory and motor system experiments (ultrafast acquisitions), the data was bandpass filtered between 0.01-0.3 Hz.

#### 4.5.2 Retinotopic mapping analysis

Retinotopic mapping analysis was performed using both conventional population receptive field (pRF) mapping ^83^ and micro-probing ^84^. In brief, these methods model the visual field locations to which a population of neurons measured within a voxel respond to as a two-dimensional Gaussian, where the center corresponds to the pRF’s position and the width to its size. In a previous study, we have implemented these techniques to map the topographic structure of the rat visual pathway ^47^.

#### 4.5.3 Layer CF model analysis

##### 4.5.3.1 Layer Definition

Histological analysis determined the cortical depth (μm) corresponding to each cortical layer, as shown in Figure 6D: LI (0–200 μm), LII/III (200–600 μm), LIV (600–900 μm), LV (900–1450 μm), and LVI (1450–2100 μm). The cortical depth for each voxel was calculated by measuring the Euclidean distance from each cortical voxel to the brain surface, performed slice by slice. The brain surface was estimated by fitting a polynomial function to several manually selected points along the boundary of the brain surface. Voxels were then assigned to cortical layers based on their calculated cortical depth and the defined layer thickness.

##### 4.5.3.2 Layer CF Model

The lCF model predicts the neural activity of a recording site (voxel) in a target layer (e.g., layer IV of V2) based on the aggregate activity in a source layer or region (e.g., the entire LP or layer II of V1). This model extends the traditional CF approach as described by^56^. Each voxel’s fMRI response is predicted using a 2D Gaussian CF model that spans across cortical layers. The key parameters of the CF model are its position and spatial spread (size) within the cortical layers. With a defined CF position and size, a predicted time-series is generated by weighting the CF model with the source BOLD time series. Optimal CF parameters are determined by minimizing the residual sum of squares between the predicted and actual time-series data. In this study, only CFs with a variance explained above 15% were retained, following criteria established in previous research ^57,85^.

#### 4.5.4 Statistical analysis

All statistical analyses were performed using MATLAB (version 2016b; Mathworks, Natick, Massachusetts, USA). Unless otherwise specified, after correction for multiple comparisons, a *p*-value of 0.05 or less was considered statistically significant. Statistical comparisons of the CF properties (size and variance explained) were performed using ANOVA with post hoc analysis Bonferroni corrected for multiple comparisons (brain area). One way t-tests were used to test whether the parabolic fits were significantly different from 0.

## Supplementary Information

**Figure S1.**
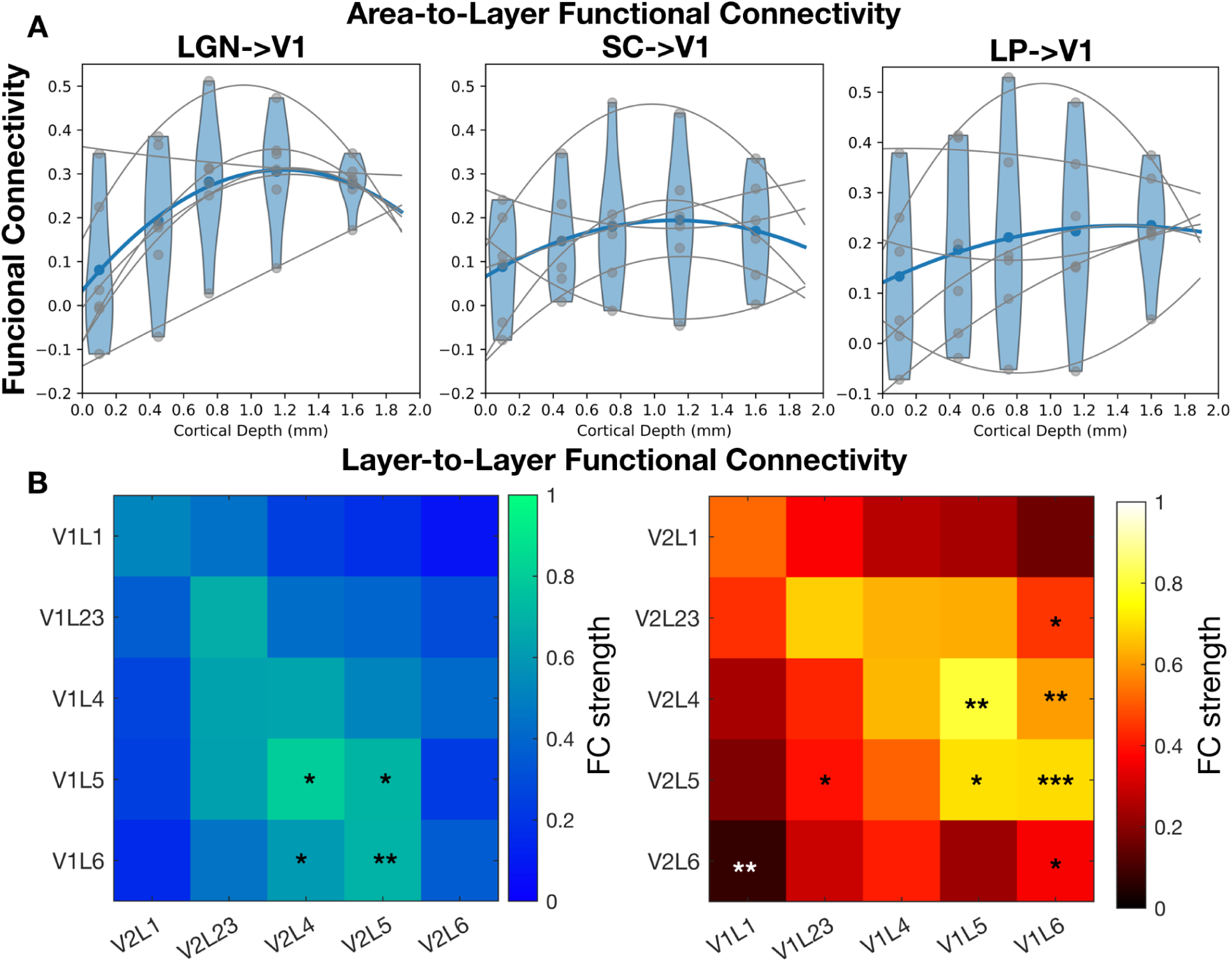
Functional connectivity Analysis. **A**: FC calculated between LGN, SC and LP, and V1 across cortical depth. The gray dots represent the data from each individual animal and gray lines the parabolic fit per animal. The blue dots and lines represent the median of all animals. **B:** FF (blue) and FB (red) connectivity maps from V1 to V2 layers obtained under spontaneous activity. The *** represents statistically significant *p*-value <0.001, ** *p*-value <0.01, and * *p*-value <0.05, calculated for the differences FC strength and lCF sizes obtained during spontaneous activity..

**Table S1.** Parabolic Coefficients obtained for the FC fit across cortical depth for the connections between LGN, LP and SC and V1.

| Functional connectivity |  |  |  |  |  |  |
| --- | --- | --- | --- | --- | --- | --- |
| Rat | LGN-V1 |  | LP-V1 |  | SC-V1 |  |
|  | Parabolic coefficient | r<sup>2</sup> | Parabolic coefficient | r<sup>2</sup> | Parabolic coefficient | r<sup>2</sup> |
| 1 | -0.05 | 0.98 | -0.05 | 0.98 | -0.05 | 0.98 |
| 2 | -0.04 | 1.00 | 0.00 | 1.00 | -0.04 | 0.95 |
| 3 | -0.03 | 0.97 | 0.01 | 0.45 | 0.00 | 0.97 |
| 4 | 0.00 | 0.66 | -0.01 | 0.76 | 0.01 | 0.75 |
| 5 | -0.03 | 0.95 | -0.01 | 0.94 | -0.02 | 0.96 |
| 6 | 0.00 | 0.99 | 0.02 | 0.89 | 0.02 | 0.93 |

**Figure S2.**
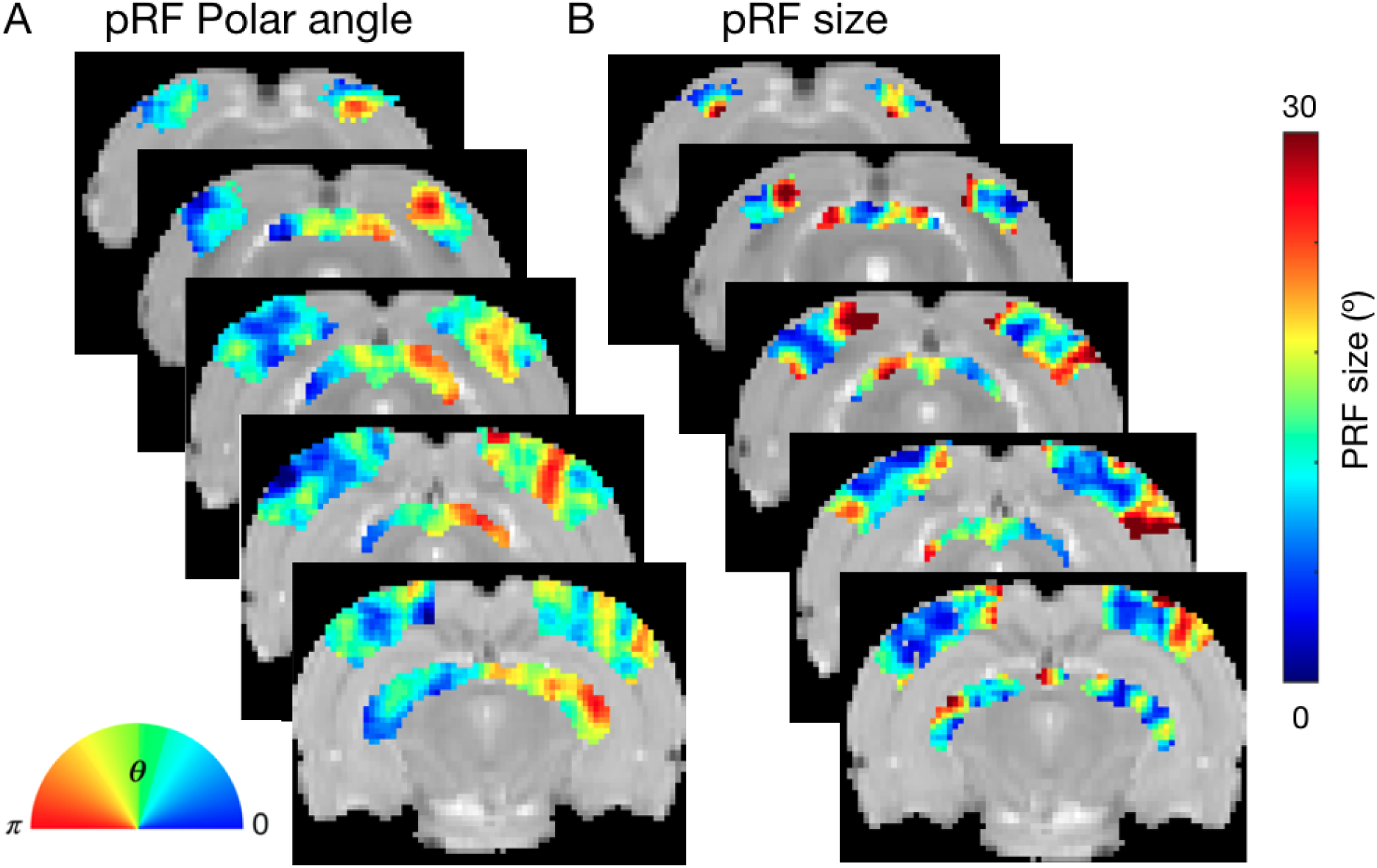
Retinotopic Mapping. **A**: Phase map of the pRF estimates projected in brain slices averaged across all animals. **B:** PRF size projected on brain slices averaged across all animals.

**Figure S3.**
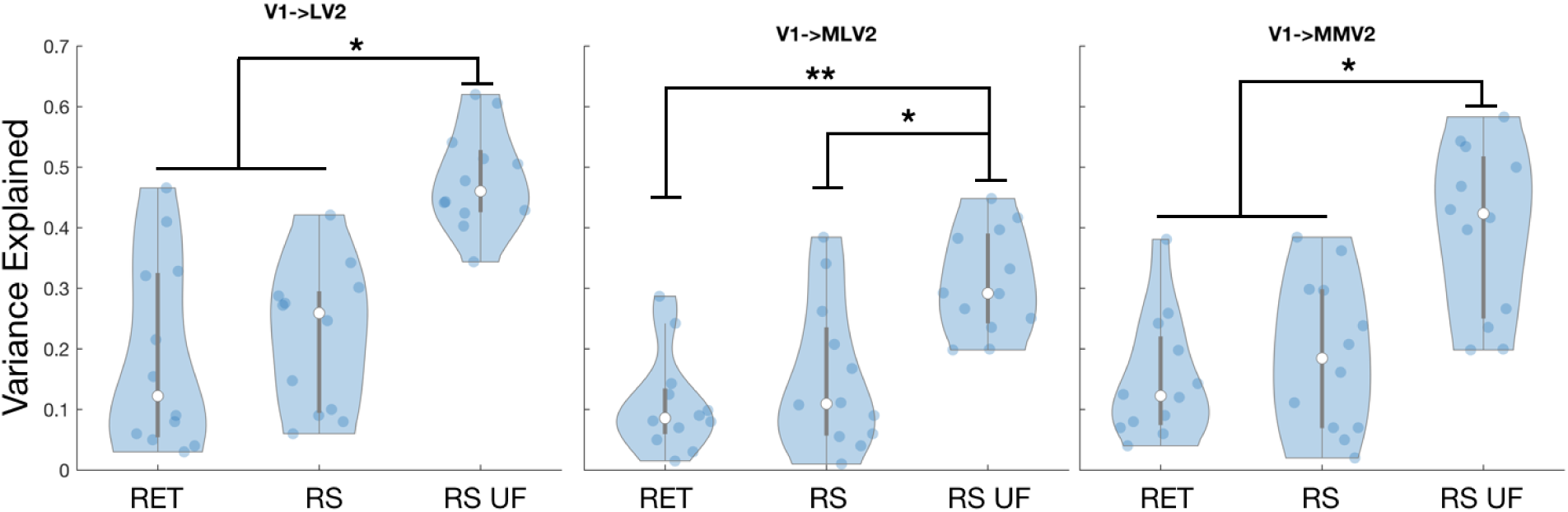
Variance explained in the lCF model obtained for visual stimulation (during retinotopic mapping) and spontaneous activity (RS) with standard temporal resolution (TR=1.5 s) and ultrafast acquisition (TR=0.05 s). The lCF model was computed from V1 to three HVA.

**Table S2.** Parabolic Coefficients obtained for the CF size fit across cortical depth for the connections LP-HVA, LP-V1, LGN-HVA and LGN-V1.

| CF size fit |  |  |  |  |  |  |  |  |
| --- | --- | --- | --- | --- | --- | --- | --- | --- |
| Rat | LP-HVA |  | LP-V1 |  | LGN-HVA |  | LGN-V1 |  |
| | Parabolic coefficient | $r^2$ | Parabolic coefficient | $r^2$ | Parabolic coefficient | $r^2$ | Parabolic coefficient | $r^2$ |
| 1 | -0.19 | 0.93 | -0.10 | 0.88 | 0.02 | 0.03 | -0.15 | 0.87 |
| 2 | -0.25 | 0.84 | 0.08 | 0.96 | -0.08 | 0.62 | -0.22 | 0.91 |
| 3 | -0.19 | 0.63 | -0.02 | 0.94 | -0.01 | 0.89 | -0.15 | 0.84 |
| 4 | -0.24 | 0.82 | -0.12 | 0.95 | 0.05 | 0.78 | -0.27 | 0.81 |
| 5 | -0.18 | 0.59 | -0.18 | 0.83 | 0.08 | 0.97 | -0.27 | 0.87 |
| 6 | -0.22 | 0.82 | -0.03 | 1.00 | -0.07 | 0.67 | -0.06 | 0.57 |

**Figure S4.**
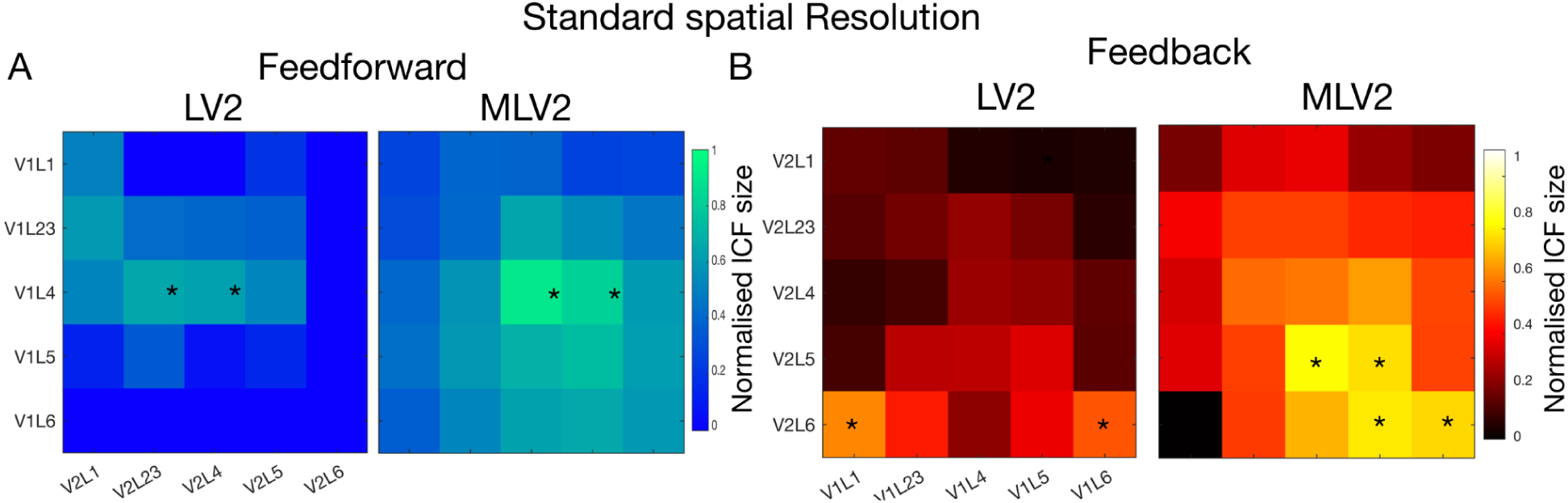
Layer-to-layer connectivity maps obtained at 50 ms TR and standard spatial resolution under spontaneous activity. **A**: FF connectivity maps from V1 to HVA (LV2 and MLV2) layers obtained under spontaneous activity. **B:** FB connectivity maps from LV2 and MLV2 to V1 layers.

**Figure S5.**
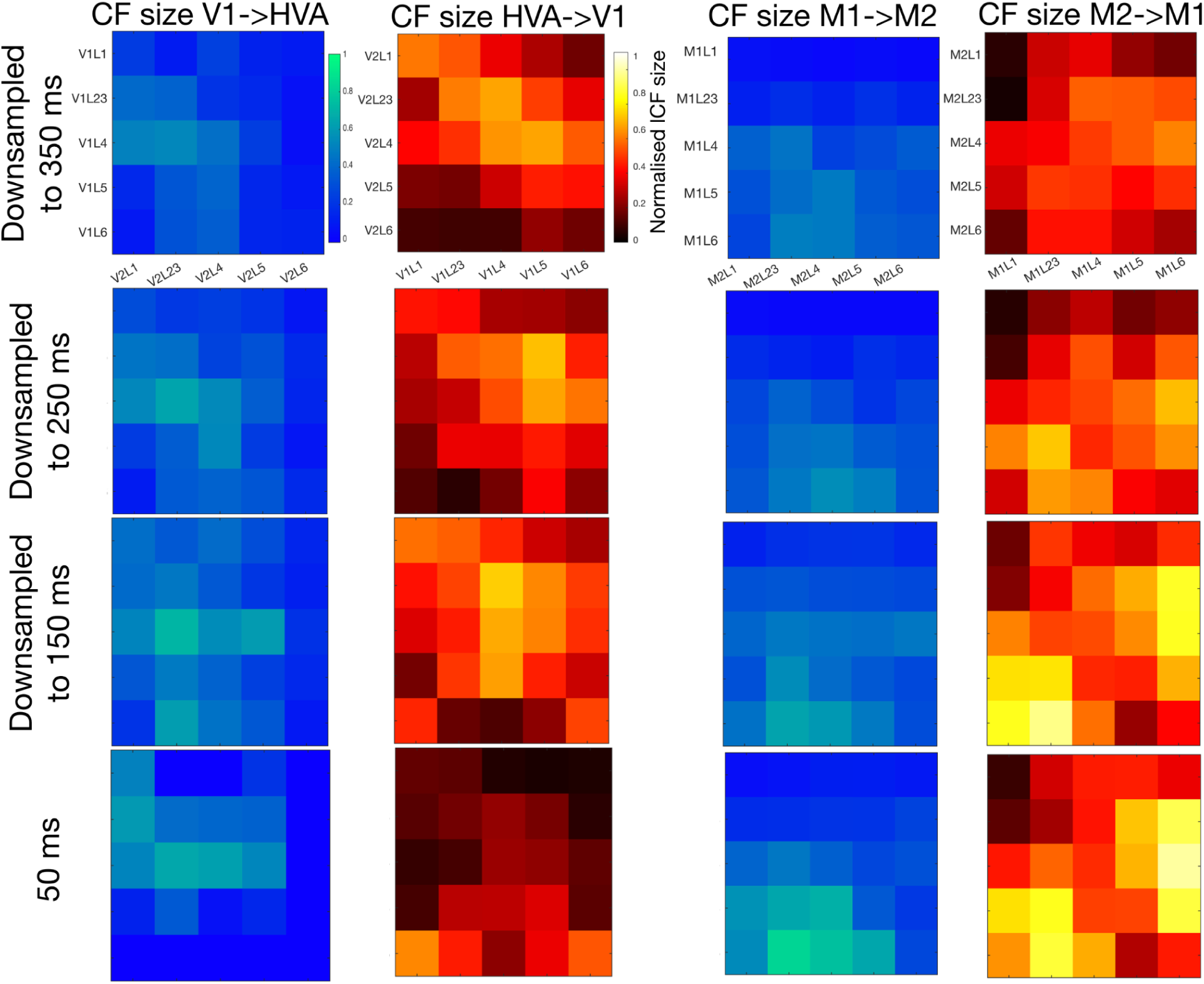
Effect of the sampling rate in the layer-to-layer connectivity maps. **Left**: FF (blue) and FB (red) layer-to-layer connectivity maps from V1 to HVA (LV2) obtained under spontaneous activity at the original sampling rate – 50 ms – and downsampled to 150 ms, 250 ms and 350 ms. **Right:** FF and FB layer-to-layer connectivity maps obtained from M1 and M2.

**Table S3.** Variance Explained of the lCF model across V1 layers obtained for healthy controls and lesioned animals for the projections LGN->V1 and LP->V1.

| Variance explained |  |  |  |  |
| --- | --- | --- | --- | --- |
|  | LGN->V1 |  | LP->V1 |  |
| V1 Layers | HC | CB | HC | CB |
| Layer I | 0.42±0.04 | 0.03±0.02 | 0.24±0.07 | 0.03±0.02 |
| Layer II/III | 0.43±0.04 | 0.04±0.02 | 0.24±0.08 | 0.03±0.02 |
| Layer IV | 0.42±0.06 | 0.04±0.02 | 0.21±0.09 | 0.03±0.02 |
| Layer V | 0.42±0.05 | 0.04±0.02 | 0.21±0.08 | 0.03±0.02 |
| Layer VI | 0.41±0.02 | 0.04±0.01 | 0.20±0.04 | 0.03±0.02 |

**Figure S6.**
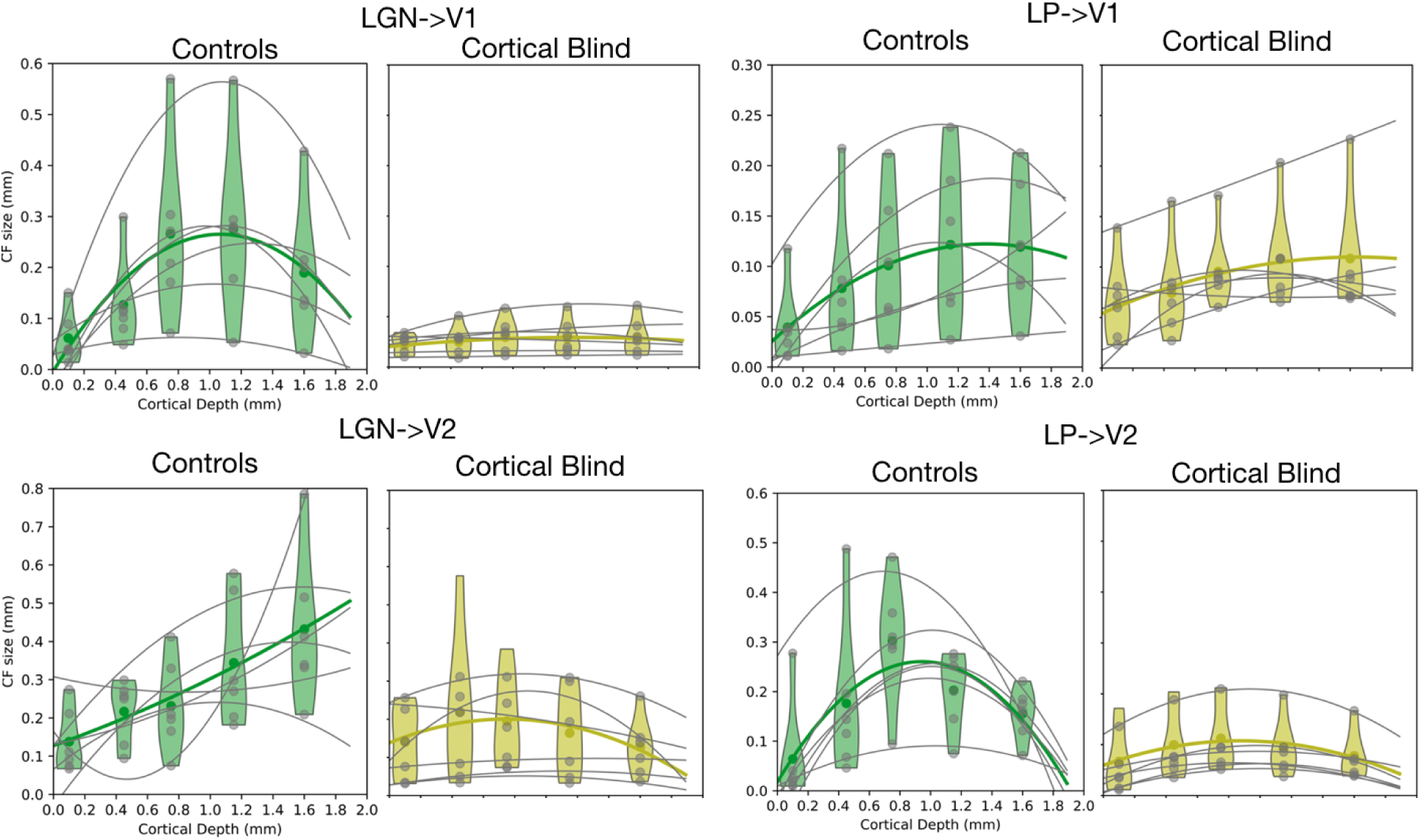
Non-Normalized lCF size. Quantitative analysis of the CF size (non-normalized) as a function of cortical depth for HC (green) and for CB animals (yellow) obtained for the connections: LGN->V1; LP->V1; LGN->HVA; LP->HVA. The gray dots represent the data from each individual animal and gray lines the parabolic fit per animal. The green or yellow dots and lines represent the median of all animals from each group.

**Figure S7.**
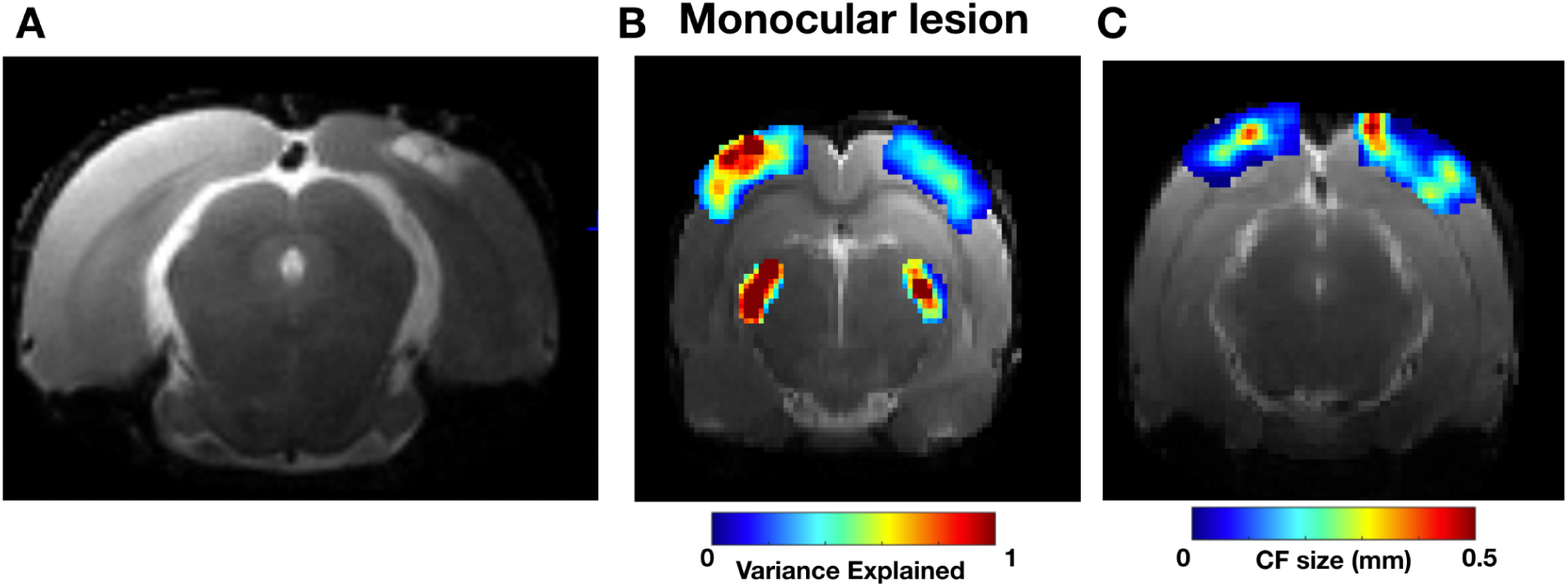
CF maps upon monocular V1 lesion. **A**: Anatomical scan of an animal with a monocular lesion on the right V1. **B and C:** CF Variance explained and CF size map using LGN as source, projected on a brain slice, respectively.

## Notes

### Competing Interest Statement

The authors have declared no competing interest.

